# How serially homologous neuroblasts produce different temporal cohorts along the *Drosophila* larval body axis

**DOI:** 10.1101/2024.11.09.622783

**Authors:** Deeptha Vasudevan, Yi-wen Wang, Hannah Carr, Elise Paniel, Sean Corcoran, Chris C. Wreden, Elaine Kushkowski, Conor Lee-Smith, Ellie S. Heckscher

## Abstract

Neural stem cells are inherently flexible, producing different lineages of neurons in different contexts; yet *in vivo*, development must be highly regulated because aberrant stem cell development leads to microcephalies, tumors, and neurodevelopmental disorders. However, the full complement of *in vivo* flexibility in neural stem cell development remains poorly understood. Within the *Drosophila* CNS, 30 types of neural stem cells or neuroblasts repeat in each segment with segment-specific differences in development. We studied NB3-3 neuroblast in each of the 14 segments where it is present. In different segments, NB3-3 produces different types of neurons in different numbers but always produces Even-skipped Lateral (EL) neurons. Prior work showed that in abdominal segments, ELs comprise two types of temporal cohorts. Temporal cohorts are small sets of neurons within a lineage that are born within a tight time window. In the abdomen, EL neurons in the early-born temporal cohort all receive mechanosensory input, whereas EL neurons in the late-born temporal cohort all receive proprioceptive input. Herein, we find that NB3-3 generates an early-born EL temporal cohort in all segments. In specific regions, NB3-3 neuroblasts produce additional types of temporal cohorts, including but not limited to the late-born EL temporal cohort. We show that variations in NB3-3 division pattern determine the number and type of temporal cohorts. Later in development, the number of neurons within a temporal cohort is refined by a variety of post-mitotic mechanisms: regulation of molecular identity markers, cell death, and differentiation status, acting in complex combinations in different segments. Post-mitotic refinement of neuron numbers occurs most often for neurons of the early-born EL temporal cohort. Further, neurons of the early-born EL temporal cohort are incorporated into different circuits, depending on segment. Together, our results show how a single neural stem cell type can flexibly generate segment-specific lineages and contribute to various circuits. We offer the following framework for understanding how a given type of neural stem cell generates diverse temporal cohorts: a stem cell type invariably generates one type of temporal cohort, regardless of location or pattern of division. For neurons in this invariant temporal cohort, post-mitotic mechanisms introduce diversity in number. The stem cell, on a segment-by-segment basis, can also produce additional types of temporal cohorts simply by altering its division pattern. This framework is valuable for motor circuit development, the development of sexual dimorphisms, nervous system evolution, and *in vitro* stem cell applications.

## Introduction

Animals have regionalized bodies requiring specialized circuits in the CNS, and yet neurogenesis starts from axially repeating stem cell pools. Over development, any given stem cell type can generate different numbers and types of neurons, depending on its location along the body axis. In the mouse spinal cord, pools of progenitors (e.g., p0, p1, p2, pMN, and p3) are found from the rostral hindbrain through the caudal spinal cord (Goulding and Lamar 2000; Francius et al. 2013). Comparisons between limb and trunk levels reveal varying numbers of motor neurons and interneurons, and greater than 10-fold differences in V1 interneuron molecular subtypes (Dasen et al., 2005; Dasen et al., 2008; Jessell, 2000; Stifani, 2014; Matsushita, 1997; Ebara et al., 2008; Sweeney et al., 2018; Francius et al., 2013). In Drosophila, serially homologous neuroblasts are found as left-right pairs in each of 16 CNS segments (O Birkholz et al., 2013; Technau et al., 2014; Kunz et al., 2012). One type of neuroblast can generate lineages of varying size with different neural types in various numbers (Birkholz et al., 2013; Technau et al., 2006; Schmid et al., 1999; Homem and Knoblich, 2012; Kraft and Urbach, 2014; Rogulja-Ortmann et al., 2008; Lacin et al., 2009; Baumgardt et al., 2014; Bossing et al. 1996; Schmidt et al. 1997; Skeath and Thor, 2003; Thor et al., 1999). However, even in the best-studied cases, the full repertoire of neurogenesis for any one stem cell type remains unknown, and an integrated understanding of which cell biological processes produce different numbers of neurons in various segments is lacking. This knowledge is crucial for understanding the inherent flexibility of stem cell development, which has implications for evolution, sex determination, and in vitro stem cell-based applications.

In this study, we used *Drosophila* CNS as a model to determine the full extent of the flexibility in neurogenesis from a single type of serially homologous stem cell. The *Drosophila* CNS is organized into an anterior hindbrain-like subesophageal zone (SEZ) and a posterior spinal cord-like nerve cord (Figure 1A). The nerve cord comprises the thorax, abdomen, and terminus, and the neurons within process stimuli from and control the movement of the head, mid-body, and tail, respectively (Diao et al., 2024). Each region is further subdivided into segments, from the anterior: the subesophageal zone (“SEZ” 3 segments), thorax (“Th” 3 segments), abdomen (“A” 7 segments), and terminus (“Te” 3 segments). Every left or right segment is called a “hemisegment.” Nearly all hemisegments contain 30 neuroblasts, with each neuroblast being a different type (Figure 1B). Herein, we study one type of neuroblast, NB3-3. NB3-3 neuroblast lineages have emerged as a powerful model for studying stem cell biology (Birkholz et al., 2013; Gunnar et al., 2008; Heckscher et al., 2015; Mark et al., 2021; Marshall and Heckscher, 2022; Touma et al., 2012; Tsuji et al., 2008; Wreden et al., 2017; Wang et al., 2022). NB3-3 neuroblasts are found as left-right pairs across 14 segments spanning from SEZ2 to Te2 (Figure 1A-B).

**Figure 1.**
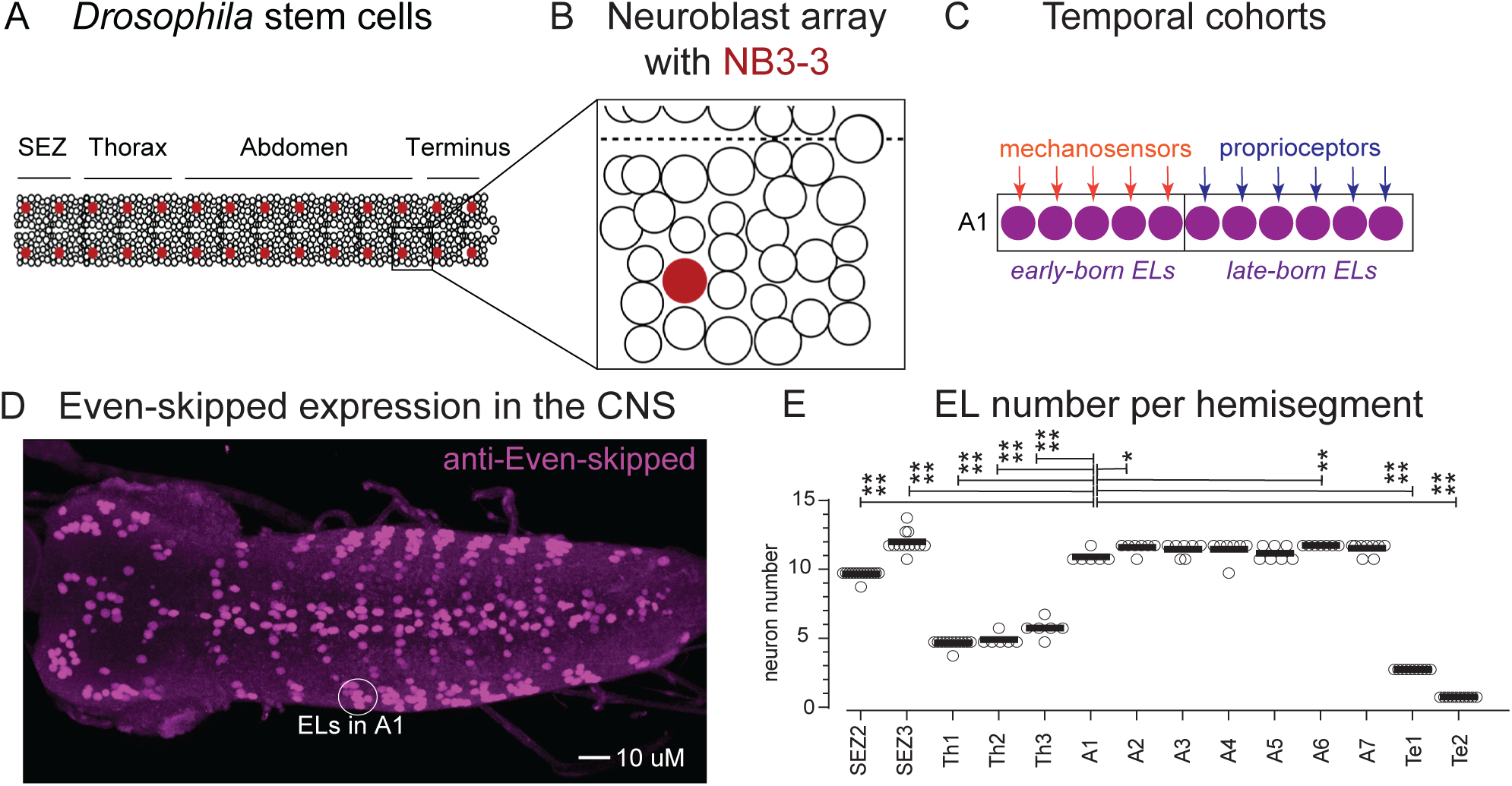
>10-fold difference in the number of ELs produced by NB3-3 in different CNS segments. (A) Illustrations of *Drosophila* neuroblasts. The developing subesophageal zone (SEZ) and nerve cord comprise a segmentally repeating set of serially homologous neuroblasts. Each circle represents one neuroblast with NB3-3 in maroon. (B) All neuroblasts within a half segment (hemisegment) are shown in the box, with NB3-3 in maroon. (C) Illustration of the inputs from various somatosensory neurons onto EL neurons in segment A1 of the nerve cord. Each circle is one Eve-expressing cell. Arrows denote direct synaptic input. Rectangles denote the groups of neurons comprising a temporal cohort. (D) Image of a wild-type *Drosophila* larval CNS stained with anti-even-skipped (Eve) antibodies. Eve Lateral neurons (ELs) are located on the lateral edges of the CNS (e.g., circle in A1). The CNS is shown in a dorsal view with the anterior left. (E) Quantification of EL number in the larval CNS. Each circle represents the EL count in a single left or right half segment (hemisegment). Black lines show the mean. All counts were compared to those in segment A1 using an ordinary one-way ANOVA **** p<0.0001; ** p<0.01; * p<0.05; only statistically significant relationships are shown.

NB3-3 neuroblasts generate lineages of neurons that include the well-studied Even-skipped (Eve)-Lateral (EL) interneurons (Bossing et al., 1996; Gunnar et al., 2008; Heckscher et al., 2015; Mark et al., 2021; Marshall and Heckscher, 2022; Touma et al., 2012; Tsuji et al., 2008; Wreden et al., 2017; Wang et al., 2022; Jay and McLean, 2019). In segment A1, the NB3-3 lineage comprises five early-born EL neurons and six late-born EL neurons (Figure 1C). Early-born and late-born ELs are examples of two different types of temporal cohorts. Temporal cohorts are lineage fragments, or small sets of neurons born from a single stem cell within a tight time window. Although first discovered in NB3-3, temporal cohorts are found in many *Drosophila* nerve cord lineages, including NB7-1, NB3-1, NB3-5, MNB, and more (Meng et al., 2019; Meng et al., 2020; Wang et al., 2022; Mark et al., 2021; Baek and Mann, 2009) as well as *Drosophila* mushroom body (Kunz et al., 2012), mouse spinal cord (Stam et al., 2012), and mouse neocortex (Gao et al., 2014). Among the NB3-3 lineages, for those in segment A1, connectome reconstruction shows that each of the five early-born ELs receives direct input from chordotonal sensory neurons, and that each of the six late-born ELs receives direct input from proprioceptive neurons (Wreden et al., 2017; Mark et al., 2021; Wang et al., 2022) (Figure 1C). Thus, the importance of temporal cohorts lies in the fact that they are lineage fragments incorporated into circuits as units.

A particularly significant gap in our understanding of temporal cohort biology is our limited understanding of diversity among temporal cohorts. There is currently one example of axial diversity in the production of temporal cohorts along the body axis: In the *Drosophila* thoracic nerve cord, NB3-3 neuroblasts enter quiescence midway through embryogenesis (Tsuji et al., 2008). Quiescence has the effect of truncating the NB3-3 lineage precisely at the border between the early-born and late-born temporal cohorts. This finding suggests that the production of temporal cohorts, rather than neuron numbers more generally, is developmentally regulated across the body axis.

In this study, we employed a combination of lineage tracing, marker gene analysis, cell death blockade, and anterograde circuit tracing to identify patterns of neurogenesis and neural development for each NB3-3 stem cell lineage in all 14 segments along the body axis. We found that the pattern of neurogenesis for the NB3-3 lineage varies along the body axis depending on the region. Nonetheless, in every segment, NB3-3 neuroblasts produce the neurons that comprise the early-born EL temporal cohort. And, depending on the specific pattern of neurogenesis, NB3-3 variably produces additional types of temporal cohorts, including, but not limited to, the late-born EL temporal cohort. After each NB3-3 neuroblast generates its specific lineage of neurons, the number, molecular identity, differentiation status, and circuit membership of these neurons are differentially tuned depending on their axial location. Tuning occurs most frequently for neurons of the early-born EL temporal cohort. Together, our data support a model in which one type of temporal cohort is invariably generated in every segment, regardless of segmental location or pattern of stem cell division. For those neurons in the invariably generated temporal cohort, post-mitotic mechanisms introduce any needed diversity in neuron number. In addition to the invariably generated temporal cohorts, additional types of temporal cohorts are generated on a segment-by-segment basis simply by altering the patterns of stem cell division. This model provides a framework for understanding the flexible development of serially homologous stem cells *in vivo*.

## Results

### Even-skipped Lateral (EL) neuron numbers differ in different segments

To begin to understand the variation in neurogenesis in NB3-3 lineages along the body axis, we determined the number of Even-skipped Lateral (EL) neurons in each segment in larvae. In late-stage embryos, EL number varies 2-fold (Heckscher et al., 2015). However, embryonic analysis quantified EL neuron number before circuit formation, which occurs in larvae, and analyzed ELs only in three of the 14 segments where NB3-3 neuroblasts are found. Herein, we identified ELs as laterally placed neurons labeled with anti-Even-skipped antibody staining and manually counted EL number (Figure 1D-E, Figure S1). In the larval terminus, we report for the first time that there is one in Te2 and three ELs in Te1. In the larval abdomen, there are 12 ELs in A2-A7 and 11 ELs in A. In the larval thorax, there are six ELs in Th3 and five in Th2 and Th1. In the two SEZ segments, Eve-expressing cells were located in four clusters. So, for segmental counts, we added EL-GAL4 driving membrane GFP as a second marker of ELs, and anti-Scr to mark the SEZ-nerve cord border (Figure S2). In SEZ3, there are 12 ELs and 10 ELs in SEZ2. In summary, by the larval stage, the numbers of ELs differ in various segments, ranging from a minimum of 1 to a maximum of 12, demonstrating segmental differences in NB3-3 lineage development.

### NB3-3 neuroblast proliferation patterns differ in the SEZ, thoracic, and abdominal segments

Neural stem cell proliferation plays a significant role in regulating the number of neurons (Homem et al., 2015), and thus, it is likely to vary across NB3-3 neuroblast lineages. Fundamentally, during development, a neural stem cell makes two decisions (collectively herein “proliferation pattern”) that can impact the number of neurons in a lineage: (1) The number of stem cell divisions (herein, division duration), and (2) whether to produce a proliferative or a post-mitotic daughter (herein, division type). In the *Drosophila* nerve cord, proliferative daughters are called Ganglion Mother Cells (GMCs). GMCs divide once to produce two cells with different fates— a Notch ON, “A” fated cell, and a Notch OFF, “B” fated cell (Truman et al., 2010; Karkavich, 2005). Generally, within the NB3-3 lineage, Notch OFF/B neurons express Eve and are ELs (Lacin et al., 2009; Lacin et al., 2014). Theoretically, a stem cell can divide any number of times from once to the entire organism’s lifetime and could produce a proliferative daughter on any division. However, it is unknown what range of proliferation patterns a population of serially homologous stem cells uses.

The proliferation patterns of NB3-3 neuroblasts are well-characterized in some but not all regions. In the thorax and abdomen, NB3-3 proliferation patterns differ from one another (Figure 2B-C). In the thorax, halfway through embryonic neurogenesis, NB3-3 neuroblasts enter a quiescent state (Tsuji et al., 2008). Before that, on each division, NB3-3 neuroblasts produce a proliferative GMC (Gunnar et al., 2016). In contrast, in the abdomen, NB3-3 neuroblasts divide throughout embryonic neurogenesis (Tsuji et al., 2008) and die before the larval stages (Baumgardt et al., 2014). Only in the first division do NB3-3 neuroblasts generate proliferative GMCs, and for the remaining divisions, NB3-3 neuroblasts directly generate Notch OFF/B neurons (Gunnar et al., 2016). We set out to determine if the SEZ and terminus used thorax-like, abdomen-like, or other proliferation patterns.

**Figure 2.**
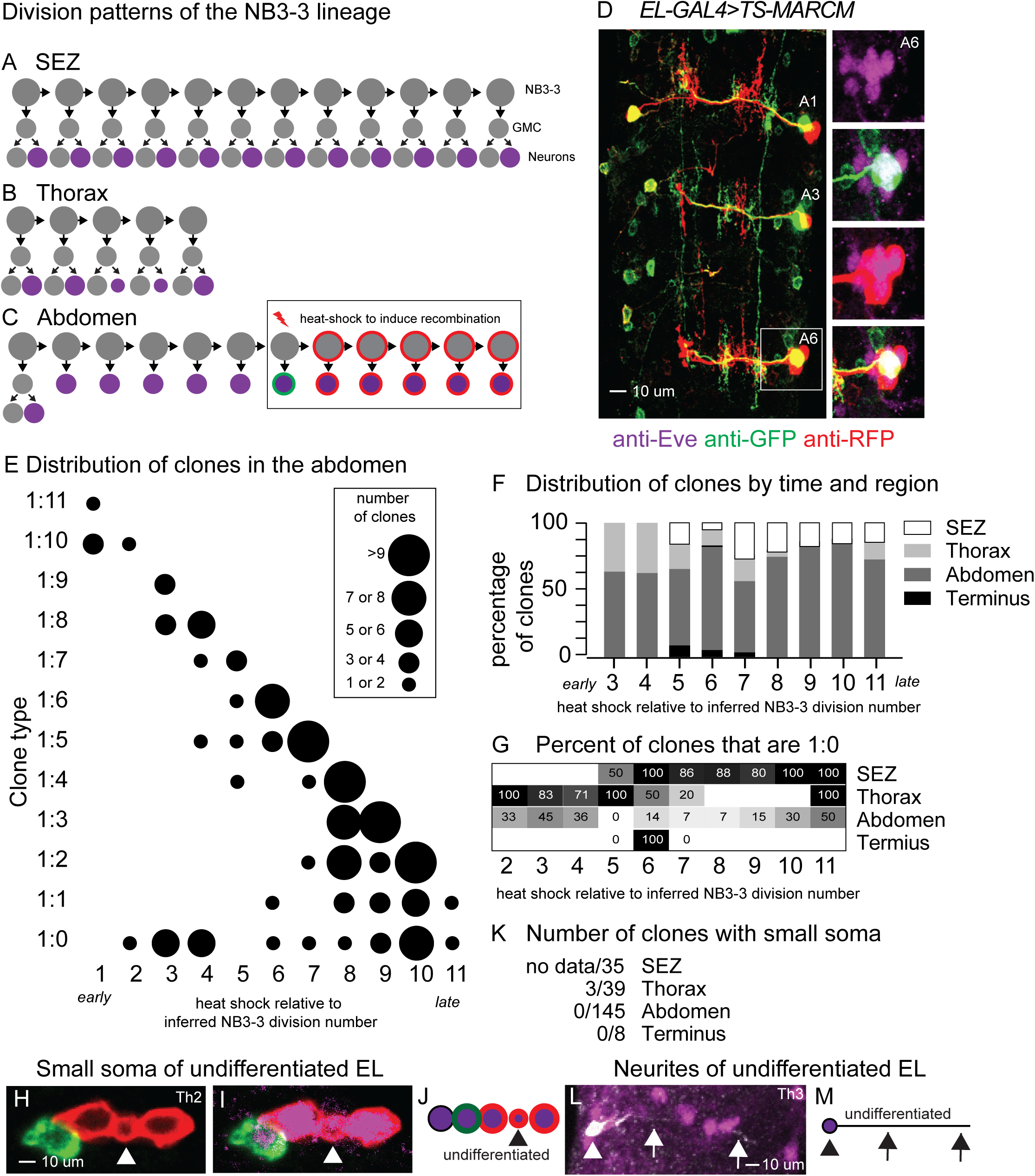
There are three distinct NB3-3 proliferation patterns along the body axis. (A-C) Illustrations of the division types of NB3-3 lineage in different regions. Each circle represents one cell, and each arrow represents a cell division. Large gray circles are NB3-3; smaller gray circles are ganglion mother cells (GMCs) or neurons. The small circle represents an undifferentiated neuron. Purple represents Eve expression. An example of the ts-MARCM induction event corresponding to the clone in segment A6R (D) is shown in the boxed region. The red lightning bolt represents the heat shock that induces a recombination event. One NB3-3 descendant is labeled with GFP (green ring) and the other with RFP (red ring). Labels are inherited by subsequent descendants (e.g., multiple red rings). Maximum abdominal clone size (RFP+ and GFP+ cells) can be used to infer heat shock time relative to NB3-3 division number, which is the X-axis in plots D-F. See methods for details. (D) Images of Twin-Spot MARCM clones. Genotype: *EL>ts-MARCM*. (A) Segments A1-A7 of the nerve cord are shown in dorsal view with anterior up. Each segment with a clone is labeled. For example, A6 refers to the CNS’s sixth abdominal hemisegment. The boxed region is enlarged in the smaller panels on the right. (E) Quantification of the distribution of clones in abdominal segments. Rows represent clone type, where #:# corresponds to the number of cells labeled in each color. For example, B shows a 1:4 clone. Columns correspond to the inferred heat shock time relative to the NB3-3 division number. Each circle indicates one or more clones were generated, with the circle size corresponding to the number of clones at each time point. Clones on the diagonal from top left to bottom right likely represent stem cell recombination events, whereas clones horizontal in the 1:0 row are likely from GMC divisions. (F, G) Quantification of the distribution of clones by time of induction and region. (F) The number of clones in each region at each heat shock time was counted and expressed as a percentage. Both early and late heat shock times generate SEZ and abdominal clones, whereas thoracic and terminal clones are produced by early heat shocks. (G) For clones at each region and time point, the number of “small (i.e., 1:0)” clones was counted and expressed as a percentage of the total number of clones in that region and time. The presence of small clones suggests the presence of a proliferative GMC daughter. In the SEZ and thorax, small clones make up most of the clones at all heat shock times. (H-J) Images and an illustration of a thoracic clone containing a small soma are indicated by the arrowhead. (K) Quantification of the distribution of clones with small (>2 microns) soma size. (L, M) Image and illustration of a thoracic clone containing a neurite with no branching (“stick-like”).

To analyze the NB3-3 neuroblast proliferation patterns, we employed an inducible lineage tracing approach, Twin-Spot Mosaic Analysis with a Repressible Cell Marker (ts-MARCM) (Yu et al., 2009). In ts-MARCM, heat shocks applied at various developmental time points deliver recombinase to random cell sets and induce a lineage trace. When the lineage trace is initiated, the daughters and all their descendants can be labeled by GFP or RFP (i.e., the twin spots) (box in Figure 2C). GFP and RFP are under UAS control and are expressed using a GAL4 line. Herein, EL-GAL4 drives the UAS reporters in EL neurons at larval stages. We combined a published ts-MARCM (Wang et al., 2022) dataset with a new one (Table S1). The differences between the datasets are (1) CNSs were imaged either at low resolution for all regions (SEZ to terminus) or higher resolution for nerve cords (thorax to terminus); (2) for the SEZ-containing dataset, inductions started at NB3-3’s 5th division. The combined data includes ∼12 different heat shock protocols, 80 CNS, and 234 clones (Table S2). Herein, “clone” refers to a group of cells labeled by one recombination event. Generally, clones were found along the entire axis, with multiple clones in each CNS segment (Figure 2D, Table S2). For analysis, we binned each CNS based on heat shock time relative to the inferred NB3-3 division number, where 1 represents the first neuroblast division and 11 represents the last division (e.g., Figure 2E). This dataset provides three new types of information: duration of division, division type, and neural differentiation status.

We can infer when NB3-3 is dividing to produce ELs because recombinase will not generate a clone if the lineage trace is induced after division stops (Figure 2F). In the SEZ and abdomen, ELs were labeled regardless of induction time. Thoracic and terminal clones were primarily generated by early induction. This difference suggests that NB3-3 neuroblasts divide to produce ELs for longer in the SEZ and abdomen than in the thorax and terminus (Figure 2A-C).

We can infer division type of the neuroblast that gave rise to the EL as follows: If a lineage trace is randomly initiated by the heat shock in a dividing GMC, the resulting clone will comprise at most two neurons (Notch ON/A and Notch OFF/B) regardless of induction time, and only the Notch OFF/B neuron will be detected by EL-GAL4. Thus, clones should appear as a single cell (herein 1:0 clones). If a lineage trace is initiated in a dividing neuroblast, the resulting clone will be large, with earlier inductions producing larger clones. Therefore, the presence of 1:0 clones suggests that NB3-3 is dividing to generate a proliferative GMC that gave rise to an EL, or that induction occurred during the final neuroblast division. We calculated the percentage of clones that were 1:0 at each heat shock time and per region (Figure 2G). For the SEZ and thorax, a majority of clones were 1:0 at nearly all induction times. For the abdomen, the percentage of 1:0 at each induction time was in the minority at all but the final induction time. For the terminus, our data was ambiguous due to the small number of clones in this region. These differences suggest that NB3-3 neuroblasts generate proliferative daughter GMCs in the SEZ and thorax on most divisions (Figure 2A-B).

We can determine the differentiation status of the EL neurons at larval stages. Differentiated neurons have a soma size of >3 microns (Heckscher et al. 2014), whereas undifferentiated neurons are smaller (∼1.5 microns). The thorax, but not other regions, possesses EL with a small soma (Figure 2B, H-K). In addition, the thoracic undifferentiated neurons also had stick-like neurites (Figure 2L, M).

So far, our ts-MARCM analyses grouped segments into regions (Figure 2A-C), however EL number varies on a segment-by-segment basis (Figure 1). Therefore, we looked for segment-by-segment differences in ts-MARCM data (Table S1). The only detectable difference was between A1 and the other abdominal segments: When both A1 and another abdominal segment were labeled in a single CNS, a majority had smaller A1 clones. These data suggest that the production of ELs by NB3-3 neuroblasts lags in A1 compared to A2-A7.

Taken together, our data show NB3-3 neuroblasts in the SEZ, thorax, and abdomen employ three distinct proliferation patterns, but they do not fully characterize the terminus. In the SEZ, NB3-3 divides for the duration of neurogenesis, producing proliferative GMCs that generate both Notch OFF/B and Notch ON/A neurons. The total SEZ lineage size should be ∼24 neurons, half being ELs (Figure 2A). In the thorax, NB3-3 divides only early in neurogenesis, again producing proliferative GMCs that generate both Notch OFF/B and Notch ON/A neurons. In this case, the lineage size should be ∼12 neurons, half being ELs (Figure 2B). In the abdomen, NB3-3 divides for the duration of embryogenesis, directly producing Notch OFF/B neurons. In this case, the lineage size should be ∼12 neurons, a majority of which are ELs (Figure 2C). Thus, the two major proliferation decisions— division type and division duration—are deployed in a combinatorial manner. Regardless of proliferation pattern, NB3-3 generates ELs early in neurogenesis in every segment. Additional neurons, including ELs that are generated later in neurogenesis and Notch ON/A neurons, are variably produced by NB3-3 in different regions, dependent on the proliferation pattern.

### In some segments, NB3-3 neuroblasts generate Notch OFF/B neurons that fail to express Even-skipped and become EL neurons

In the prior section, we used ts-MARCM to assay the proliferation patterns of NB3-3 in various segments. However, these experiments had four limitations: (1) NB3-3 development in the terminus was not fully characterized; This is important as prior publications disagree on the possibility of proliferative daughter GMCs in the NB3-3 lineage in the terminus (Birkholz et al. 2013; Gunnar et al. 2016). (2) The viability of the NB3-3 neuroblasts was not determined. This is important because prior reports raise the possibility that neuroblast death in mid-neurogenesis could limit the number of ELs in the terminus (Birkholz et al. 2013). (3) Our conclusions so far are mainly on the regional rather than segmental level (Figure 2); yet we observe differences in EL number segment by segment (Figure 1). (4) ELs but not any other NB3-3 progeny were tracked. However, we need to track all NB3-3 progeny because one possible mechanism limiting the number of ELs is that some Notch OFF/B neurons fail to express Eve, and therefore, to become ELs. Indeed, connectome analysis showed that the first-born, Notch OFF/B neuron in A1 remains undifferentiated and fails to express Eve (Wang et al., 2022).

We employed a second lineage tracing approach, the <u>G</u>AL4 <u>T</u>echnique for <u>R</u>eal-time <u>A</u>nd <u>C</u>lonal <u>E</u>xpression (G-TRACE) (Evans et al., 2009), in which transient GAL4 expression is converted into permanent nuclear-localized GFP labeling of descendant cells. We use the G-TRACE with Eagle-GAL4, which is transiently expressed in four classes of neuroblasts and a subset of their progeny (Higashijima et al., 1996). Among labeled cells, the NB3-3 lineage can be identified based on position in all segments except SEZ2. In addition to G-TRACE, we stained with anti-ElaV (a pan-neuronal marker), anti-Hey (a Notch activity marker), and anti-Eve (the EL marker) (Figure 3A). With this combination, on a segment-by-segment basis, we can identify four cell types in the NB3-3 lineage: the NB3-3 neuroblast itself (GFP[+]), Notch ON/A neurons (GFP, ElaV, Hey[+]), Notch OFF/B neurons without Eve expression (GFP, ElaV[+]) and Notch OFF/B neurons with Eve expression, that is EL neurons (GFP, ElaV, Eve[+]) (Figure 3B-C).

**Figure 3.**
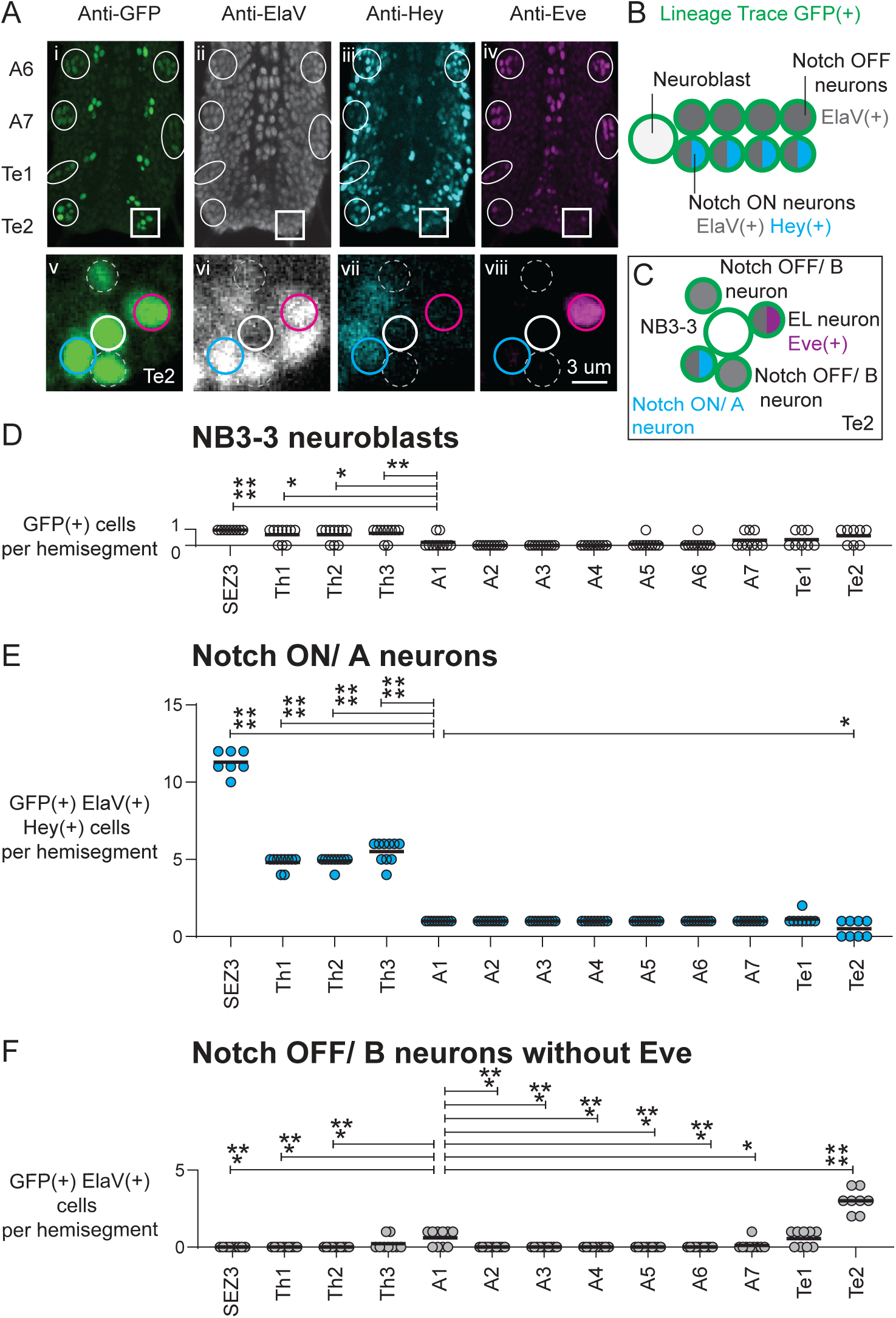
In A1, Te1, and Te2, some NB3-3 neural progeny fails to express Even-skipped. (A) Images of a lineage tracing experiment. Segments A6-Te2 of the *Drosophila* embryonic nerve cord (stage 17) are shown in ventral view with dorsal up. The tissue was stained with anti-GFP(i,v), anti-ElaV(ii, vi), anti-Hey(iii, vii), and anti-Eve (iv, viii). The image is a thick z-section including all NB3-3 neuronal progeny ovals and box) and a subset of other labeled neurons. In this example, the NB3-3 lineage trace failed in the Te1 right segment. The Te2 right segment is enlarged in v-viii. (v-viii) A single confocal slice shows NB3-3 (white circle), an NB3-3A neuron (cyan circle), an EL neuron (magenta circle), and two NB3-3B neurons that do not express Eve (dashed circle). Genotype: Eagle>G-TRACE. (B, C) Illustrations of lineage markers. Each circle is a single neuron. Colors represent gene expression. The green circle is the GFP lineage trace. Gray is neural ElaV expression. Cyan is Hey (Notch reporter) expressing. Magenta is Eve expression. (B) The illustration represents a typical relationship between a neuroblast and its A- and B- hemilineage neural progeny. (C) The illustration represents data shown in Av-viii. (D-F) Quantifications of the distribution of NB3-3 neuronal types. (D) At the end of embryogenesis, NB3-3 neuroblasts are usually found in the SEZ and thorax, variably found in the terminus, and rarely in the abdomen. (E) Many NB3-3A (Notch ON) neurons are found in SEZ and thorax, but not the abdomen or terminus. (F) NB3-3B neurons without Eve are found in segment A1, Te1, and Te2. Each circle represents the cell count in a single hemisegment. Bars show averages. ANOVA comparisons are versus A1. **** = p<0.0001, *** = p<0.001, ** = p<0.01; * = <0.05. Only statistically significant relationships are shown. Genotype: *Eagle>G-TRACE*.

We began asking which NB3-3 neuroblasts die by counting NB3-3 neuroblasts per segment at the end of embryogenesis. We reasoned that if NB3-3 was dead, we would not detect it in our lineage trace data. NB3-3 is present in the majority of thoracic and SEZ segments, and ∼half of the terminal segments, but not in the abdominal segments (Figure 3D). These data suggest that in the abdomen but not elsewhere NB3-3 dies, and rule out neuroblast cell death as the cellular mechanism limiting EL in the terminus.

Next, we assayed the division type of NB3-3 in all segments by counting the number of NB3-3 Notch ON/A neurons in each segment (Figure 3E), with the idea that Notch ON/A neurons are only produced by proliferative GMCs. In the abdomen, each segment possesses one Notch ON/A neuron, and terminal segments possess one or no Notch ON/A neurons. These data suggest that NB3-3 only produces one GMC in the abdominal and terminal segments. In contrast, SEZ3 possesses 12 Notch ON/A neurons. Th1 and Th2 possess five Notch ON/A neurons, and Th3 possesses six. In these segments, the number of Notch ON/A neurons matches exactly the number of EL neurons (Figure 1). Taken together with ts-MARCM data (Figure 2), these data suggest that in the thorax and SEZ, on every division, NB3-3 produces a GMC that divides to make one Notch OFF/B (i.e., EL neuron), and one Notch ON/A (non-EL) neuron.

Finally, we looked for segments in which failure to express Eve in Notch OFF/B neurons limited EL numbers, counting the number of Eve(-) Notch OFF/B neurons (Figure 3F). In our assay, in A1, a majority of segments had one Notch OFF/B neuron that failed to label with Eve. In contrast, in A2-A7, all Notch OFF/B neurons were labeled with Eve (i.e., none failed to label with anti-Eve). Combined with other data (Figures 1-2), this suggests NB3-3 stem cells in abdominal segments A1-A7 exhibit the same proliferation pattern, but, in segment A1 only, the first-born Notch OFF/B neuron fails to differentiate and express Eve. In the terminus, a subset of Notch OFF/B neurons fail to express Eve: one in Te1 and three in Te2. In the SEZ and thorax, all Notch OFF/B neurons are labeled by Eve. We conclude that a new mechanism for controlling the number of ELs is the failure to express Eve in Notch OFF/B neurons. This mechanism is prominent in, but not exclusive to, the terminus because it is also observed in segment A1.

### Programmed cell death limits the number of EL and non-EL neurons in terminal segments only

Cell death is a candidate for regulating the number of neurons in various segments along the body axis. In the CNS of mice, C. elegans, and Drosophila, ∼50% of embryonically born neurons die during normal development. In *Drosophila* embryos, blocking cell death in neuroblasts and neurons increases NB3-3 lineage in Te2 size from six to 10 neurons (Birkholz et al., 2013). However, which cells die in the terminus, and the extent to which cell death occurs in the NB3-3 lineage in other segments, remain unknown.

We began by quantifying EL numbers in posterior segments A7, Te1, and Te2, starting at mid-neurogenesis. Between mid-neurogenesis and larval stages, EL numbers doubled in A7 and were reduced by half in Te1 and Te2 (Figure 4A-D, F). To determine if the reduction in ELs was due to programmed cell death (PCD), we used two approaches. We removed RHG genes required for neural PCD using the Df(3L)H99 mutant (Figure 4E), or as a second approach, we inhibited apoptosis using a pan-neural driver to drive the p35 Caspase inhibitor (herein Elav>p35, Figure 4G-H). The results were similar: EL number increases in the terminus from three to five in Te1 and from one to two in Te2 (Figure 4F, I). Additionally, in the Elav>p35 background, we counted the EL number in A7 and other segments in more anterior regions (SEZ3, Th2, A2) and found the number unchanged (Figure 4I).

**Figure 4.**
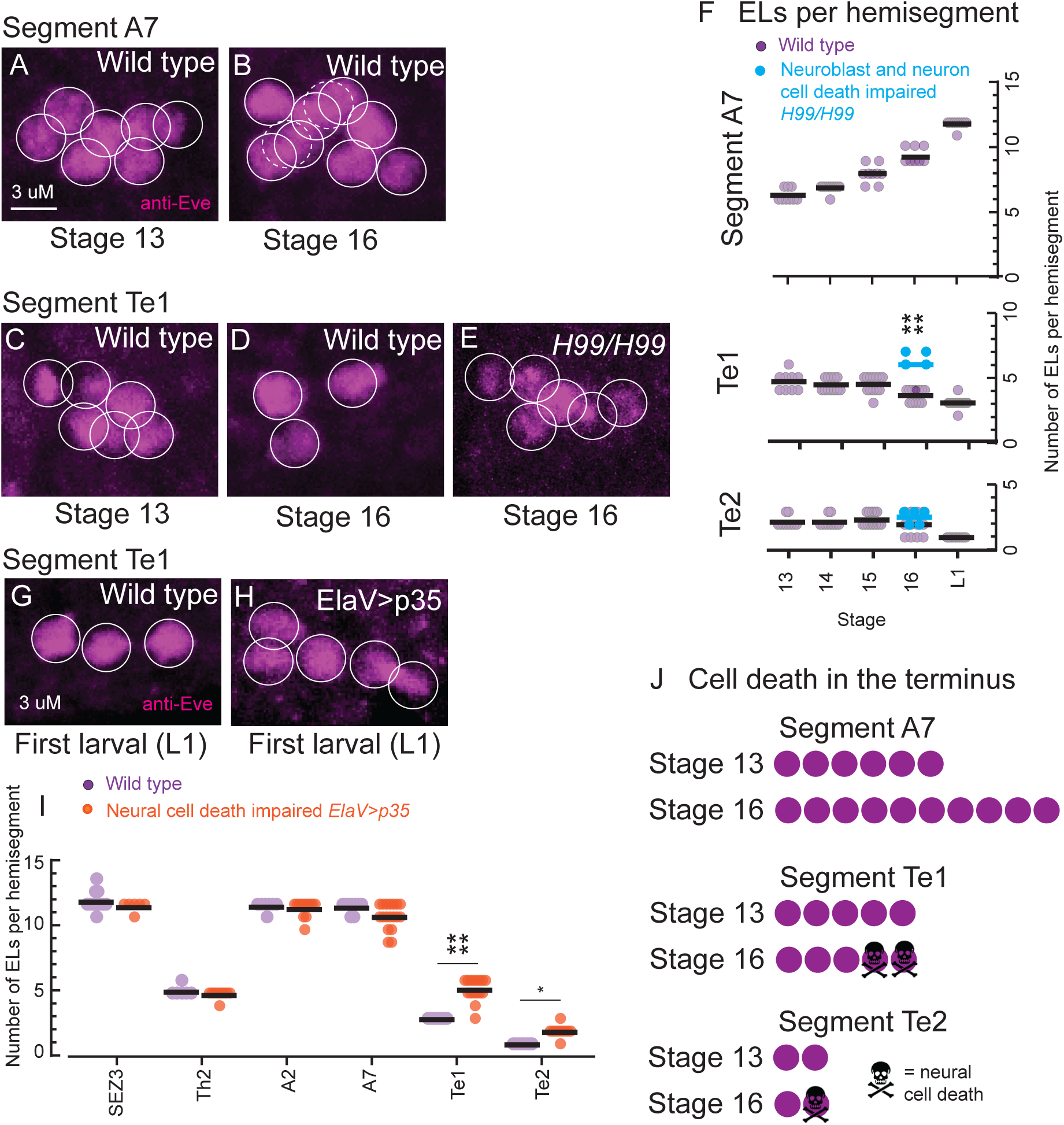
A subset of EL neurons undergoes programmed cell death in the terminus but not elsewhere. (A-H) Images of nerve cords stained with anti-Eve show ELs in posterior nerve cord segments in wild type and when cell death is blocked. Each circle denotes one EL, with dashed circles in (B) indicating the ELs obscured by the image stack. (A-D, G) Wild-type tissue shows baseline EL number at each stage. (E) H99/H99 mutants block neuroblast and neural cell death in embryos. (H) ElaV>p35 blocks cell death in neurons. (F, I) Quantification of EL number in various segments. Each circle represents the EL count in a single right or left half (hemisegment). Bars show averages. (F) EL numbers for three segments—A7, Te1, and Te2—over development. (I) EL numbers at larval stages for segments spanning the body axis are shown for wild type and when neural cell death is blocked. T-test compares paired data (genotypes in each segment). **** = p<0.0001, * = <0.05. Genotypes: light purple is wild type; cyan is *Df(3L)H99/Df(3L)H99*, and orange is *ElaV>p35*. ANOVA comparisons to values in Figure 1E. Only statistically significant relationships are shown. (J) Illustration summarizing cell death of ELs. Each circle represents an EL.

To assess the dynamic production and loss of non-EL NB3-3 progeny, we counted neuron numbers using the G-TRACE dataset. We compared the neuron numbers in each segment between two late embryonic stages (i.e., stages 15 and 17). Notch ON/A neuron numbers are constant in nearly all segments (Table S3), except for SEZ3, where the number of Notch ON/A neurons increased (Table S3), suggesting that Notch ON/A neurons live. In contrast, Eve(-) Notch OFF/B neurons are constant in number in all segments <u>except</u> the terminus, where they decrease from five to three in Te2 and from one to zero in Te1 (Table S4).

We conclude that in the terminus, but not elsewhere, ∼50% of EL neurons are pruned by programmed cell death after birth (Figure 4J). Furthermore, both Eve(+) (i.e., ELs) and Eve(-) Notch OFF/B neurons of the NB3-3 lineage are killed.

### All segments have ELs with early-born molecular identities, but only a subset have ELs with late-born molecular identities

In segment A1, NB3-3 neuroblasts generate two types of temporal cohorts—early-born ELs and late-born ELs (Wreden et al., 2017; Mark et al., 2021; Wang et al., 2022). Temporal cohorts have been identified using three different methods; In all cases, abdominal segments were the focus. (1) Initially, temporal cohorts were identified as a developmental unit based on the shared expression of one molecular marker, 11F02-GAL4, that is expressed in late-born, but not early-born ELs (Figure 5A-B) (Wreden et al., 2017). (2) Connectome reconstructions with inferred lineage and birth time information have found sets of neurons with similar synaptic partnerships and called these temporal cohorts (Mark et al., 2021; Wang et al., 2022). (3) Herein, we used lineage tracing to identify temporal cohorts based on the timing of neuroblast divisions (Figure 2). More specifically, in this study, we found that in all segments, NB3-3 produces ELs early in neurogenesis. In the SEZ and abdomen, NB3-3 produces ELs later in neurogenesis. Thus, in addition to identifying temporal cohorts via a third method, our lineage tracing experiments suggest that the number and types of temporal cohorts produced by NB3-3 neuroblasts vary along the body axis.

**Figure 5.**
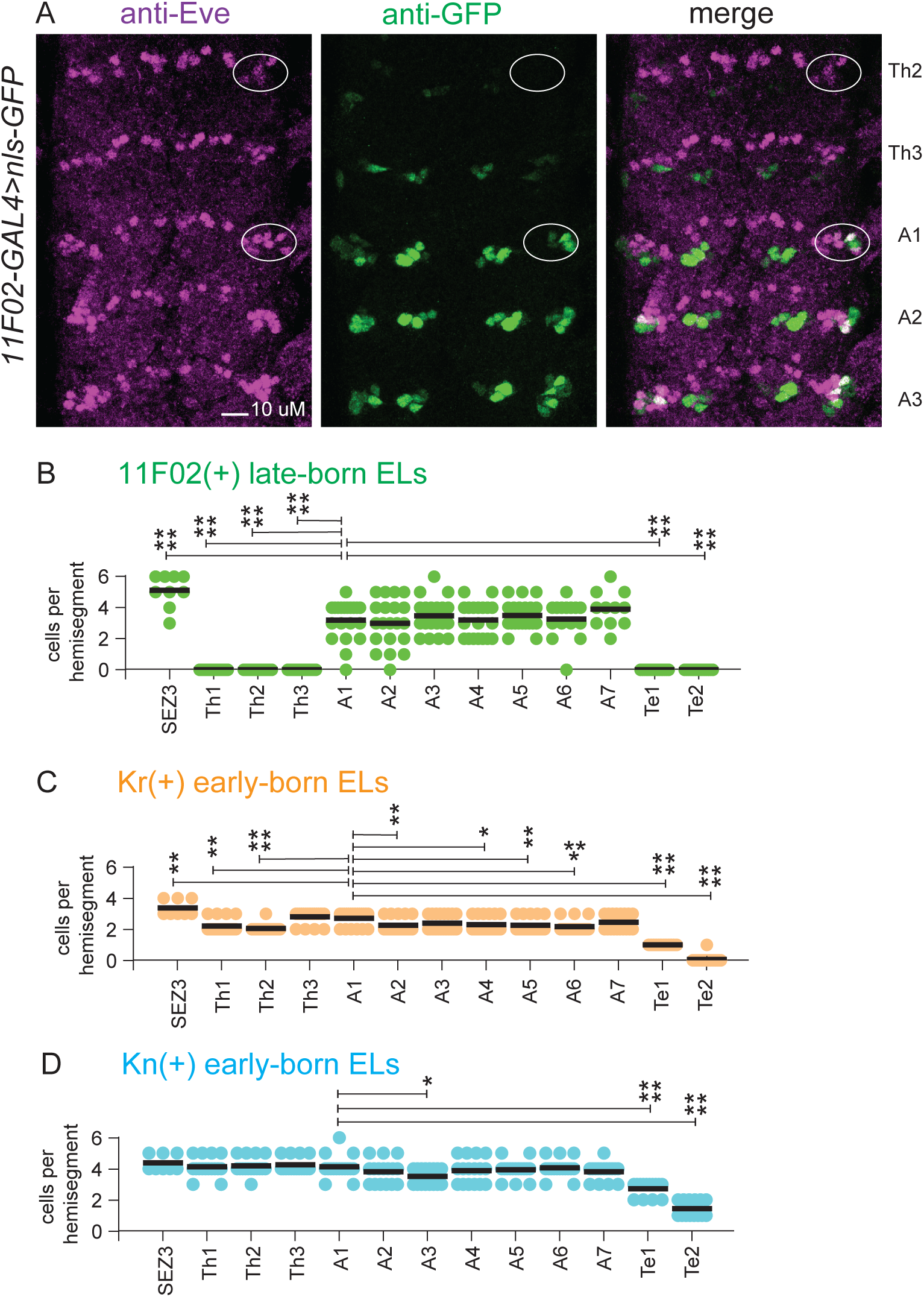
ELs with early-born molecular identity markers are found in all segments, but ELS with late-born molecular identity markers are found in the SEZ and abdomen only. (A) Images of the expression pattern of the late-born EL identity marker, 11F02, in segments Th2 to A3 of a late-stage embryonic nerve cord. The nerve cord is shown in a ventral view with the anterior up. The EL neurons on the right side of the Th2 and A1 segments are circled in each image. Genotype: *11F02>nls-GFP* (B-D) Quantification of the number of ELs expressing the given marker. Each circle represents the cell count in a hemisegment. The average is shown as a black bar. ANOVAs for multiple comparisons to counts in A1. **** = p<0.0001, *** = p<0.001, ** = p<0.01; * = <0.05; only significant differences are shown.

To test the idea that the number and types of temporal cohorts produced by NB3-3 neuroblasts vary along the body, we characterized the distribution of EL temporal cohorts staining for molecular markers. Based on our lineage tracing, one prediction is that the early-born EL temporal cohort should be found in every segment. A well-characterized marker that labels a subset of early-born ELs is the transcription factor Kruppel (Kr)(Tsuji, Hasegawa, and Isshiki 2008; Wreden et al. 2017). Kr labels between 2-4 ELs in the SEZ, thorax, and abdomen, and one EL Te1, but none in Te2 (Figure 5C). To confirm that the EL in Te2 indeed belongs to the early-born temporal cohort, we used an additional marker of early-born ELs with broader expression, the transcription factor and Knot/Collier (Kn) (Figure 5D) (Demilly et al., 2011). Generally, four ELs are labeled by Kn across SEZ, thoracic, and abdominal segments; In the terminal segments, all ELs are labeled by Kn. Thus, as predicted by lineage tracing, NB3-3 generates a temporal cohort of early-born ELs in every segment.

Based on our lineage tracing, a second prediction is that the late-born EL temporal cohort is found only in the SEZ and abdomen. We crossed 11F02-GAL4 to a nuclear-localized GFP (UAS-nls-GFP), stained late-stage embryos with anti-GFP and anti-Eve (Figure 5A-B). ELs with 11F02 labeling are found in abdominal segments A1-A7. In the SEZ, ELs are organized into four clusters of ∼5-6 neurons each (Figure S2A), each of which could correspond to a temporal cohort. Consistent with this possibility, we found that one of the two clusters of ELs in SEZ3 is labeled by 11F02 (Figures 5B and S2D). Expression of 11F02 in SEZ2 was ambiguous. ELs in the thorax and terminus do not express 11F02. Thus, as predicted by lineage tracing, NB3-3 generates a temporal cohort of late-born ELs in the SEZ and abdomen, but not thorax or terminus.

### Early-born temporal cohorts can be mapped into different circuits depending on the axial region

Temporal cohorts are fragments of lineages that can be mapped into specific circuits (Meng and Heckscher 2021). In this section, we address the following questions: (1) To what extent do ELs in different segments receive synaptic input from chordotonal (CHO) sensory neurons? In segment A1, all early-born ELs get direct synaptic input from CHO sensory neurons that respond to mechanical vibrational stimuli (Wang et al., 2022). Both early-born ELs and CHO sensory neurons are found in every nerve cord segment (Figure 6A, Bodmer et al., 1987). (2) To what extent do ELs in different segments receive synaptic input from proprioceptors? In segment A1, all late-born ELs get direct synaptic input from various proprioceptive sensory neurons (Wang et al., 2022). Proprioceptors, like dorsal bi-polar dendrite (DBD), are present from segments Th2 to Te1 (Figure 6A-B), but late-born ELs are found in abdominal (and SEZ) segments only.

**Figure 6.**
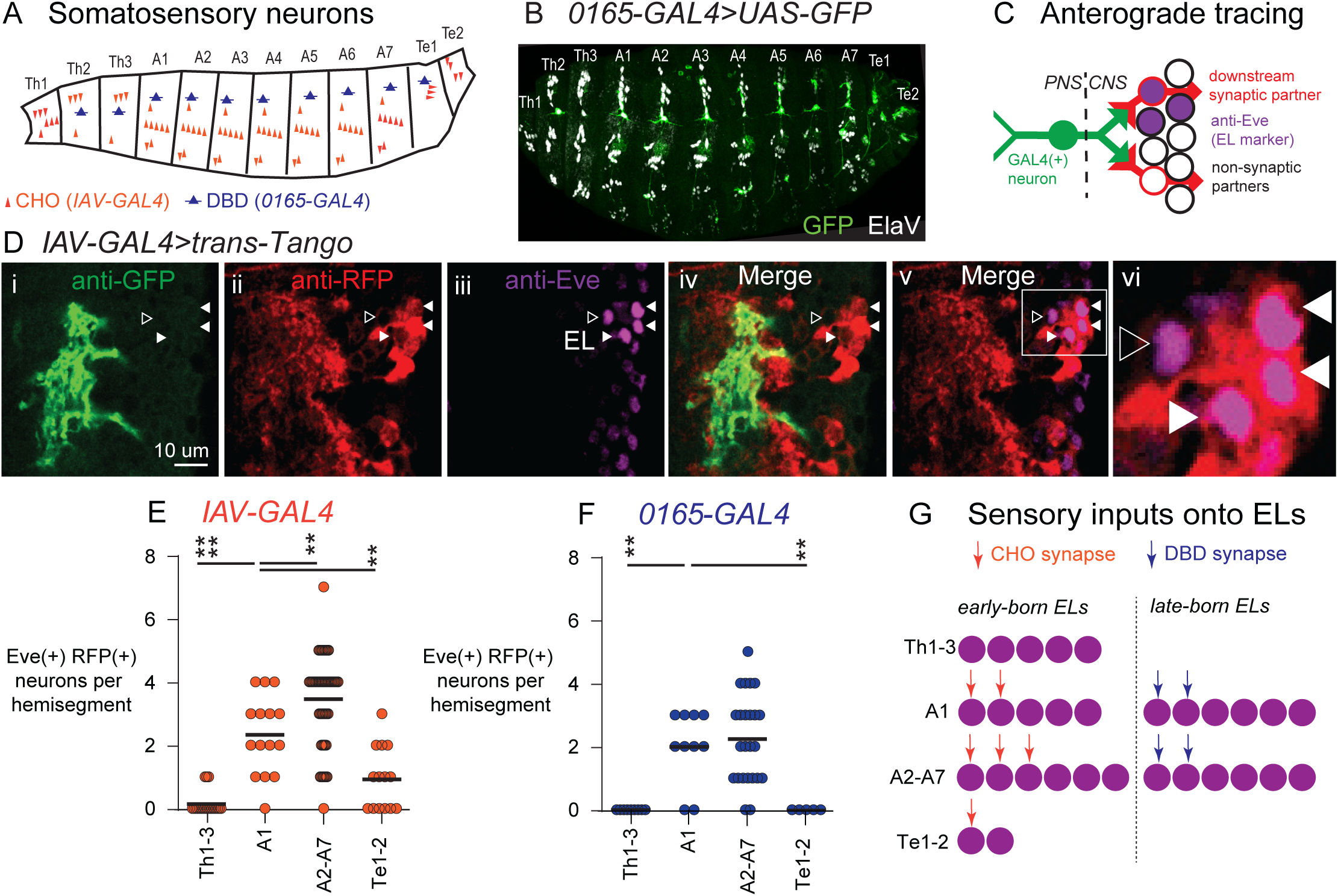
Abdominal and terminal ELs receive input from chordotonal and bipolar dendrite somatosensory neurons, but ELs in the thorax do not. (A) Illustrations of somatosensory sensory neurons in Drosophila larvae. Chordotonals (CHO) are vibration sensory neurons, shown in orange, labeled by *IAV-GAL4*. Dorsal bipolar dendrite neurons (DBD) are stretch sensors, shown in dark blue, labeled by *0165-GAL4*. (B) Image of a *Drosophila* embryo at stage 17 showing GFP expression in DBD neurons, co-labeled with ElaV (white). Anterior is to the left, and dorsal is up. The image is a maximum intensity Z-stack projection. Genotype: *0165-GAL4>UAS-GFP* (C) Illustration of the trans-Tango anterograde labeling system. trans-Tango labels neurons expressing GAL4 in green. Neurons that receive direct synaptic input from GAL4(+) neurons become RFP(+) and are labeled in red. Non-labeled neurons are represented as hollow black circles. Purple circles represent anti-Eve staining of nuclei of EL interneurons. (D) Image of an example trans-Tango experiment. A single image slice of a hemisegment at A4 of an L3 CNS is shown in dorsal view with anterior up, midline to the left, and scale bar of 10 microns. Arrowheads point to ELs visible in this focal plane. Solid arrows are ELs labeled by RFP, and hollow arrows are unlabeled by RFP. (Di) shows CHO sensory neuron termini in the CNS, (Dii) shows the membrane of neurons that form synapses with CHO sensory neurons. (Diii) shows anti-Eve staining. (Div) shows the close apposition of green and red signals, consistent with their synaptic partnership. (Dv) shows that ELs receive synapses from CHOs. (Dvi) is an enlargement of (Dv). Genotype: *IAV-GAL4*>*trans-Tango* (E, F) Quantification of sensory neuron inputs onto ELs by region. Each circle represents the cell count in a hemisegment. The average is shown as a black bar. Only significant differences are shown in ANOVAs for multiple comparisons to counts in A1 **** = p<0.0001, ** = p<0.01. Genotypes: *IAV-GAL4*>*Trans-Tango*; *0165-GAL4*>*Trans-Tango* (G) Illustration summarizing the number of inputs to ELs by region.

To identify somatosensory inputs onto ELs, we used the trans-Tango anterograde circuit tracing technique. In trans-Tango, the starting (GAL4-expressing) neurons are labeled with GFP, and their downstream synaptic partners are labeled with RFP (Figure 6C, D). To identify downstream targets of CHOs and DBDs, we used *IAV-GAL4* and *0165-GAL4* (Marshall and Heckscher, 2022), respectively. We counted the number of RFP[+] ELs in each segment to identify those that received input. First, we calibrated trans-Tango for use in larval *Drosophila*, focusing on segment A1, where connectome data are available (Wang et al., 2022). In the connectome, of the five early-born ELs in A1, three are strongly connected to CHOs (>15 synapses), two are weakly connected (<15 synapses), and late-born ELs are unconnected (0 synapses). In trans-Tango, in A1, ∼2 ELs are labeled using *IAV-GAL4* (Figure 6E). Similarly, in the connectome of the six late-born ELs in A1, three are strongly connected to proprioceptors, three are weakly connected, and the early-born ELs are unconnected. In trans-Tango, in A1, ∼2 ELs are labeled using *0165-GAL4* (Figure 6F). These data suggest that the trans-Tango method identifies ELs strongly synaptically (>15 synapses) connected to somatosensory neurons.

Trans-Tango tracing from *IAV-GAL4* labeled two ELs in A1, three in A2-A7, one in Te1-Te2, and none in Th1-3 (Figure 6E). Trans-Tango tracing from *0165-GAL4* labeled numerically identical ELs in abdominal segments and failed to label ELs in thoracic or terminal segments (Figure 6F).

Our results are consistent with the following model (Figure 6G): Although CHO sensory neurons and early-born ELs are present in every segment, they are only synaptically coupled to each other in a subset of segments (i.e., within the abdomen and terminus), and the number of neurons receiving strong input varies. DBD sensory neurons are present in segments Th2-Te1, but only late-born ELs in the abdomen get their input. When synapses do occur, they are numerically similar from segment to segment. Although we did not examine inputs onto ELs in the SEZ, SEZ ELs are likely mapped to different circuits because the SEZ does not receive projections from CHO or DBD body wall somatosensory neurons.

## Discussion

In this study, we determined the full extent of the flexibility in neurogenesis from a single type of serially homologous stem cell *in vivo*. We performed this work in the context of temporal cohort biology, revealing how temporal cohorts are diversified along the body axis. To do so, we tracked NB3-3 neuroblast lineages in 14 segments of the *Drosophila* CNS, using a combination of lineage tracing (Figures 2-3), marker gene analysis (Figures 1, 5), cell death blockade (Figure 4), and anterograde circuit tracing (Figure 5). NB3-3 neuroblasts invariably produce the neurons that comprise the early-born EL temporal cohort in every segment (Figure 4). Depending on the region, the proliferation pattern for the NB3-3 lineage varies (Figures 2-3) variably producing additional neurons that comprise different types of temporal cohorts, including, but not limited to, late-born EL temporal cohorts (Figures 3, 5). After each NB3-3 neuroblast generates its specific lineage of neurons, neuron numbers within a temporal cohort are trimmed using various post-mitotic mechanisms, including the induction of neural programmed cell death (Figure 4), establishment of molecular identity (Figure 3, 5), and regulation of differentiation status (Figures 2, 3). Post-mitotic mechanisms selectively affect neurons within the early-born EL temporal cohort. Further, circuit membership can be modulated for neurons in early-born EL, but not late-born EL temporal cohorts (Figure 5). Together, our data support the following model: Serially homologous stem cells invariably produce one type of temporal cohort regardless of location or stem cell proliferation pattern. For this type of temporal cohort, post-mitotic mechanisms introduce diversity in number for neurons, and further variation at the level of circuit wiring can also occur. Various additional types of temporal cohorts are generated simply by altering the neuroblast proliferation patterns. This model provides a framework for understanding the flexible development of serially homologous stem cells *in vivo*.

### NB3-3 lineage development varies in different segments of the CNS

Prior studies provided us with a fragmentary understanding of how NB3-3 lineage development varies across segments, which is typical for studies of the development of most *Drosophila* neuroblast lineages (Birkholz et al., 2013; Gunnar et al., 2008; Heckscher et al., 2015; Mark et al., 2021; Marshall and Heckscher, 2022; Touma et al., 2012; Tsuji et al., 2008; Wreden et al., 2017; Wang et al., 2022). Herein, we provide a single-cell-level analysis of NB3-3 lineage development in every segment where NB3-3 is present, offering new comprehensive insight.

Combining our data, the following picture emerges (represented in Figure 7): In SEZ2, NB3-3 divides 10 times (Figure 2F), producing a GMC that divides to produce Notch ON/ A and Notch OFF/ B neuronal pairs on each division (Figure 2G, 3E), with all Notch OFF/ B neurons differentiating and expressing Eve (Figure 1F, 3F, 4I); NB3-3 is alive at the end of embryogenesis (Figure 3D). In SEZ3, the development of the NB3-3 lineage is similar to that of SEZ2, but it divides only 12 times. In Th1-Th3, development of the NB3-3 lineage is similar to the SEZ with the following differences: (1) NB3-3 stops dividing after the fifth or sixth division (Figure 2F); (2) A small subset of Notch OFF/ B express Eve but fail to further differentiate at larval stages (Figure 2H-M). In A2-A7, NB3-3 divides 12 times, producing in the first division a GMC that generates a Notch ON/ A and Notch OFF/ B neuron pair (Figure 2F, G) and in the remaining divisions, producing only Notch OFF/ B neurons. All Notch OFF/ B neurons in A2-A7 express Eve and differentiate (Figure 1F, 3F, 4I), and NB3-3 dies at the end of neurogenesis (Figure 3D). In A1, development of the NB3-3 lineages is similar to that of A2-A7, except that the first Notch OFF/ B neuron fails to express Eve (Figure 3F). In Te1-Te2, the development of the NB3-3 lineage is similar to A2-A7, with the following differences: (1) NB3-3 is often alive at the end of embryogenesis. (2) NB3-3 divides six or seven times. (3) The Notch OFF/ B neurons can: express Eve and survive, express Eve and die, lack Eve and live, or lack Eve and die.

**Figure 7:**
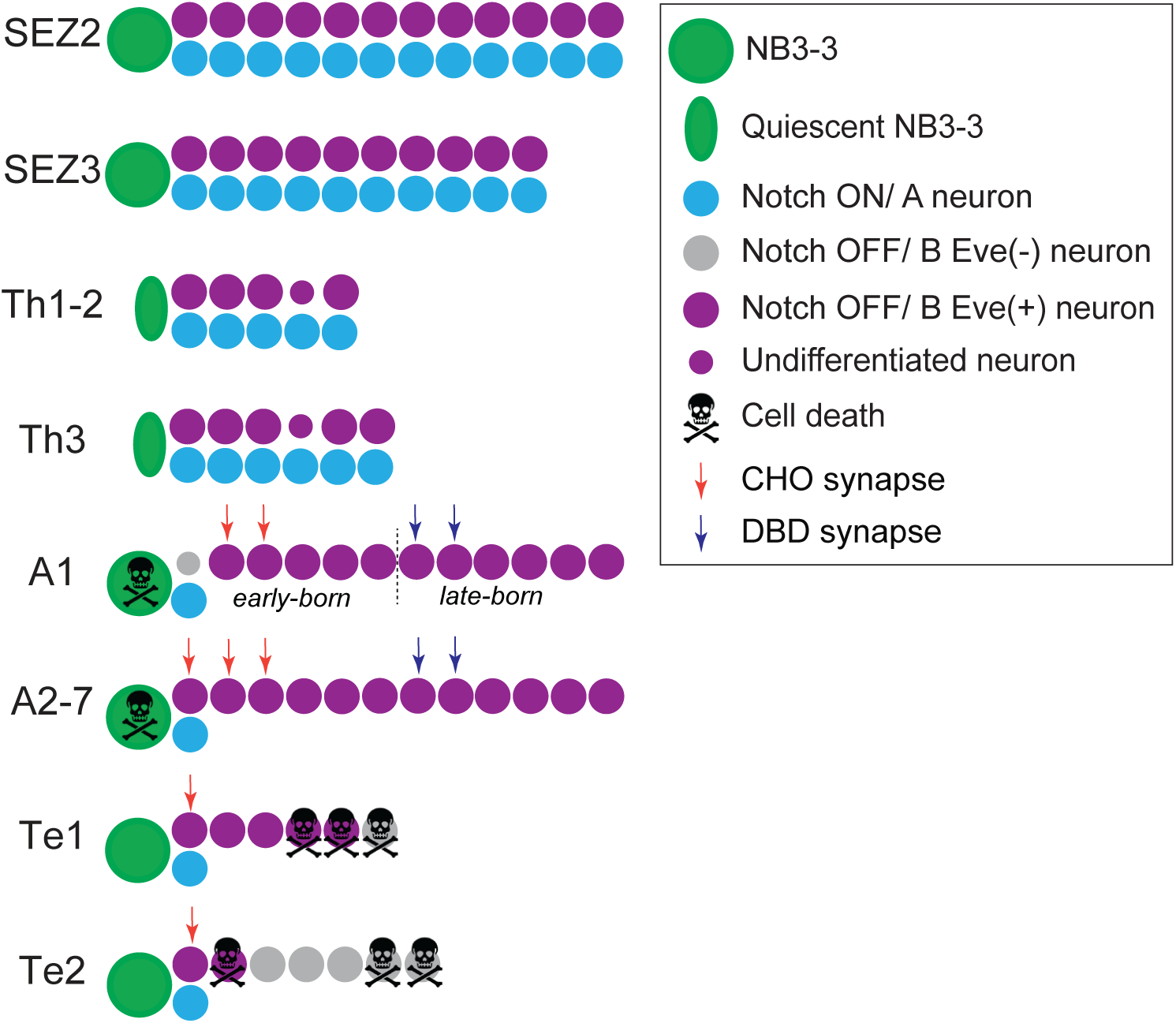
Summary of NB3-3 lineage progression in the Drosophila nerve cord from anterior to posterior segments. Illustration of neurons produced by NB3-3 in different regions of the body axis. The box shows figure legends. In segment A1, the boundary between the early-born and late-born temporal cohort is depicted with a dotted line. Every circle represents one neuron and is quantitatively accurate with the number of NB3-3 progeny produced per hemisegment.

To put these results into a larger phylogenetic context, we note how the NB3-3 lineage displays trends observed in other phyla. (1) The highest neuron numbers are found in the anterior CNS (Cobeta et al., 2017). Indeed, the distribution of neuron numbers in the NB3-3 lineage follows this trend. (2) In the mammalian spinal cord, more neurons are present in regions that control limbs (Francius et al., 2013). Analogously, EL numbers do not smoothly taper from anterior to posterior; instead, the largest number of ELs is found in two non-adjacent regions, SEZ and the abdomen. (3) In the mouse spinal cord, ∼10-fold differences in molecular subtypes for V1 neurons (Sweeney et al., 2018). In *Drosophila*, NB3-3 neuroblasts show differences in EL number, depending on region, with similar fold changes, suggesting this trait is shared across phyla. Thus, for this study and future research, NB3-3 development now offers a uniquely tractable, detailed, and comprehensive model for studying how stem cells flexibly produce neurons.

### Serially homologous NB3-3 neuroblasts use a small subset of proliferation patterns to generate different numbers and types of temporal cohorts in different regions

Stem cell proliferation is tightly regulated, as defects can cause microcephaly or megaloencephaly, tumors, or neurodevelopmental disorders (Liszewska and Jaworski, 2018; Homem et al., 2016; Swartling et al., 2015; Ossola and Kalebic, 2022). The two stem cell proliferation parameters that must be regulated are the duration of stem cell division and whether the stem cell produces a proliferative daughter. In theory, any stem cell could divide throughout the organism’s lifetime, producing proliferative daughters at every division, such that there is an extremely large number of possible proliferation patterns. It is unknown for any given stem cell type, which of the available proliferation patterns it uses— every combination? One? A subset? And, if it is a subset, what “rules” dictate what proliferation patterns are used? Our data raises the possibility that serially homologous neuroblasts use a small subset of possible proliferation patterns, and that what is being regulated is the number of temporal cohorts produced, rather than neuron number.

From the literature, we considered two models for how serially homologous neuroblasts might select proliferation patterns. Under a <u>null model</u>, the durations and types of proliferation would vary stochastically across segments, resulting in a continuous and unstructured distribution of neuron numbers (Llorca et al., 2019). In a <u>unitary production model</u>, based on the vertebrate neocortex, there is a fixed neurogenic output of ∼8–9 neurons per progenitor (Gao et al., 2014). However, our data support a third model, a <u>multimodal production mode</u>l. In a multimodal model, serially homologous neuroblasts generate different numbers of neurons depending on the segment. In the case of NB3-3, neuron number varies in increments of ∼5-7 neurons, corresponding to a unit called a temporal cohort. More specifically, depending on the specific pattern of neurogenesis, NB3-3 variably produces between one and four temporal cohorts. In all segments, NB3-3 generates one early-born EL temporal cohort. In the terminus, this is the only temporal cohort produced. In the abdomen and thorax, NB3-3 produces two temporal cohorts, but the type of temporal cohort produced other than early-born ELs varies. In the abdomen, NB3-3 neuroblasts additionally generate late-born ELs, whereas in the thorax, NB3-3 additionally generates early-born Notch ON/A neurons. In the SEZ, NB3-3 produces four types of temporal cohorts— early-born and late-born EL temporal cohorts, as well as early-born and late-born Notch ON/ A neurons.

### EL numbers are modulated in early-born EL temporal cohorts

An unexpected observation was that EL numbers are modulated in the early-born EL temporal cohorts but not late-born EL temporal cohorts. We had expected the opposite based on the following reasoning: A fundamental challenge for animals that develop from neural stem cells is that not all neurons, circuits, and resulting perceptions or actions can be produced simultaneously, so animals prioritize the production of specific neurons. One example is motor neurons, which are among the first-born neurons in most organisms and are essential for all movement (Meng et al., 2020). The motor neuron example highlights the assumption that early-born neurons are of fundamental importance and less likely to vary compared to later-born neurons. Reinforcing this idea is work from zebrafish, showing that fast behaviors or those involving gross movements emerge first, as well as the neurons critical for those behaviors (Fetcho, Higashijima, and McLean 2008; Fetcho and McLean 2010). In externally developing animals (e.g., zebrafish), following such a developmental strategy would allow for fast escape behaviors before refined movements, which likely provides a fitness benefit. Our data suggest that the opposite may be true in some cases, as we find more variation in early-born EL temporal cohorts (Figure 7). We can only speculate about why early-born temporal cohorts may be more diverse. Two speculative possibilities are: (1) In analogy to a gene duplication and divergence model, adding late-born temporal cohorts may allow for the divergence of early-born temporal cohorts. (2) Early-born ELs may be a basal neuron type and thus have had more time to evolve.

### Apoptosis is one of several mechanisms to regulate the number of EL neurons

Neuronal programmed cell death (PCD) occurs across phyla and is often essential for proper nervous system function (Sulston & Horvitz, 1977; Ellis & Horvitz, 1986; Sulston et al., 1983; Oppenheim, 1991; Yamaguchi & Miura, 2015; Buss et al., 2006). Neural PCD occurs in C. elegans, the *Drosophila* visual system and nerve cord, as well as vertebrate neocortex and spinal cord (Okado et al., 1984; Homma et al., 1994; Frade and Barde, 1999; Sulston and Horvitz, 1977; Sulston et al., 1983; Avery and Horvitz, 1987; Bello et al., 2003; Tan et al., 2011; Abrams et al., 1993; Fan and Bergmann, 2014; Blanquie et al., 2017; Southwell et al., 2012). Moreover, PCD can occur in a segment-specific manner (Bello et al., 2003; Miguel-Aliaga and Thor, 2004; Birkholz et al., 2013; White et al., 1994). Together, this has led to a model in which the neural number is determined by making an excess and killing a precise number. Our study adds complexity to this simple model. We find that in addition to PCD (Figure 5), other non-PCD mechanisms refine EL numbers (i.e., either failure to differentiate or express Eve or both) (Figures 2-3). Because we can analyze ELs at a single-cell resolution (Figure 7), we can quantify and compare the effects of PCD and non-PCD mechanisms, finding that PCD is more restricted both in terms of region and the number of neurons affected. Regionally, neuronal PCD in the NB3-3 lineage occurs only in the terminus, whereas non-PCD mechanisms reduce EL numbers in the thorax, abdomen, and terminus. Numerically, 10 neurons undergo PCD, whereas 20 use non-PCD mechanisms (four do both). These data suggest that non-PCD mechanisms— including the regulation of marker gene expression and neural differentiation status— are overlooked, yet potentially widespread mechanisms that contribute to the sculpting of post-mitotic neuron number alongside apoptosis.

Why would the CNS generate neurons that neither differentiate nor die? While the role of undifferentiated neurons remains unclear, one clue is that undifferentiated neurons are mostly present in the early-born EL cohort, not the late-born EL cohorts. Consistent with their early birth, undifferentiated neurons might play a role early in neurogenesis, such as providing trophic or physical support for later-born neurons. A second, not mutually exclusive idea is that during pupation, undifferentiated neurons may form axons and dendrites to populate adult circuits.

### In various regions, complex and distinctive combinations of cell biological mechanisms act at both the stem cell and neuron level to sculpt the NB3-3 lineage

A prerequisite for understanding the developmental logic of the CNS is an integrated understanding of the cell biological processes that diversify homologous lineages. Studies have elucidated the molecular mechanisms underlying the axial regulation of one cell biological processes at time, including neural stem cell proliferation, specification of motor neuron identity, and regulation of neurotransmitter type (Bahrampour et al., 2019; Cobeta, et al., 2017; Gouti and Gavalas, 2008; Seifert, 2015; Homem et al., 2015; Tsuji et al., 2008; Berger et al., 2005; Jungbluth et al., 1999; Hobert, 2021; Jung et al., 2010; Dasen and Jessell, 2009; Dasen et al., 2005; Kessel, 1994; Feng et al., 2022; Lints and Emmons, 1999; Estacio-Gómez and Díaz-Benjumea, 2014; Suska et al., 2011; Feng et al., 2020). Yet, we do not understand how these mechanisms are integrated.

Hox genes are major molecular regulators of patterning along the body axis, in both vertebrate and invertebrate nervous systems (Pearson et al., 2005; Philippidou and Dasen, 2013; Estacio-Gómez et al., 2013). Different segments of the body express distinct combinations of Hox genes (Dasen et al., 2005; Tsuji et al., 2008; Cobeta et al., 2017). Generally, in *Drosophila* embryos, the pattern of Hox gene expression, starting from the anterior, is as follows: Pb, lab, Dfd, Scr, Antp, Ubx, abdA, AbdB, which has two isoforms, AbdB.m and AbdB.r (Joshi et al., 2022; Birkholz et al., 2013; Diao et al., 2024). Correlating these expression domains with the various NB3-3 serially homologous lineage progression models shown in Figure 7 leads to a simple model by which one Hox expression domain promotes one type of lineage development. This simple model remains to be tested.

In the Hox literature, a Hox-specificity paradox exists— despite their region-specific functions, different Hox proteins frequently bind to identical DNA motifs *in vivo*, leading to widespread questions about how binding specificity is achieved at the level of downstream gene regulation (Slattery et al., 2011; Crocker et al., 2015; Kribelhauer et al., 2020). This paradox underscores the importance of understanding not only which Hox genes are expressed, but also the combinatorial cell biological processes occurring at both the stem cell and neuronal levels in different regions. Our data indicate developmental degeneracy in NB3-3 lineage progression, as there is no one-to-one correlation between NB3-3 lineage progression and the repertoire of cell biological decisions made by the NB3-3 lineage in a given region. Instead, the NB3-3 lineage deploys a complex combination of converging and diverging cell biological mechanisms. By converging, for example, Eve is not expressed in Notch OFF/ B neurons in the Ubx(+) A1, AbdB.m(+) Te1, and AbdB.r(+) Te2 segments. Such convergence correlates with the Hox-specificity paradox, in that different Hox genes likely have similar molecular targets. By diverging, for example, all Notch OFF/ B neurons are viable in the Ubx(+) A1 segment, but not in the AbdB.m(+) and AbdB.r(+) terminal segments. So, Ubx should not regulate the same repertoire of cell biological processes compared to AbdB, even if they both likely promote programmed cell death in the NB3-3 lineage. Thus, other factors must provide the specificity, allowing for divergence. Chromatin context, Hox co-factors, or post-transcriptional regulation (Slattery et al., 2011; Mazzoni et al., 2013; Johnston and Desplan, 2008) are all likely to play important roles.

### Conclusion and Outlook

In summary, we have identified a combination of cell biological processes that act in both stem cells and circuits, generating different numbers of neurons from one type of serially homologous stem cell. This provides a new framework for understanding variation in stem cell development. Because these insights are fundamental, they have wide-ranging implications across fields— for the evolution of behavior, development of sexually dimorphic circuitries, motor systems biology, the production of diverse organoids, etiology of disease, and regenerative medicine.

## Methods

### *Drosophila* Strains and Culture

Fly stocks were maintained at 25°C on a standard cornmeal molasses medium. Experimental crosses were set up at the indicated temperatures. The following fly lines were used:

| Name | Genotype | Location | Source |
| --- | --- | --- | --- |
| Wild type | w<sup>1118</sup> | X | (Marshall and Heckscher 2022)<br>(Heckscher et al. 2015) |
| H99 | Df(3L)H99 | 3 | BL #1576 |
| p35 | UAS-p35 | 2 | BL #5072 |
| Elav-GAL4 | Elav-GAL4 | X | BL #5144 |
| G-TRACE | P{w[+mC]=UAS-RedStinger}4,<br>P{w[+mC]=UAS-FLP.D}JD1,<br>P{w[+mC]=Ubi-<br>p63E(FRT.STOP)Stinger}9F6 | 2 | Evans et al., 2009;<br>BL #28280 |
| EL-GAL4 | EL-GAL4 | 3 | Heckscher et al.,<br>2015 |
| Twin Spot MARCM | UAS-rCD2.RFP.UAS-GFPi/CyO;<br>TM3/TM6B | 2 | BL #56184 |
| Twin Spot MARCM | UAS-mCD8.GFP.UAS-rCD2i/CyO; TM3/TM6B | 2 | BL #56185 |
| 11F02-GAL4 | P{R11F02-Gal4 (w+)}attP2 | 3 | BL#49828 |
| UAS-nls-GFP | P{UAS-GFP::lacZ.nls}30.1, (w+) | 3 | BL# 6452 |
| UAS-CD4-GFP | y[1] w[*]; P{w[+mC]=UAS-CD4-<br>tdGFP}8M2 | 2 | BL #35839 |
| TransTango | y[1] w[*] P{y[+t7.7]<br>w[+mC]=UAS-myrGFP.QUAS-<br>mtdTomato-3xHA}su(Hw)attP8;<br>P{y[+t7.7] w[+mC]=trans-<br>Tango}attP40 | 1,2 | Talay et al., 2017;<br>BL# 77124 |
| IAV-GAL4 | IAV-GAL4 | 2 | BL#36360 |
| 0165-GAL4 | 0165-GAL4 | 3 | Marshall et al., 2022 |

Embryos and early-stage larvae were used. At these developmental stages, flies have no distinguishing sexual characteristics. Thus, all experiments were conducted in a manner that was blind to sex.

### Tissue preparation

Four tissue preparations were used: (1) Late-stage whole mount embryos, in which antibodies can still penetrate the cuticle; (2) late-stage dissected embryos, in which the neuromuscular tissue and cuticle are dissected away from the CNS, allowing for superb immuno-labeling and visualization of the entire axial nerve cord from SEZ to terminus; (3) isolated larval CNSs, in which the CNS is removed from other larval tissue so that antibodies reach the CNS’s. For all preparations, standard methods were used for fixation in fresh 3.7% formaldehyde (Sigma-Aldrich, St. Louis, MO). Embryos were staged for imaging based on standard morphologic criteria.

### Immunohistochemistry

Tissue was washed three times with phosphate-buffered saline containing 0.1% Triton X-100 (PBT), blocked 1 hr in PBT containing 2% normal goat serum (NBT), and stained with primary antibody in PBT at 4°C overnight. After washing, samples were stained with secondary antibody at room temperature for 1 hr, washed again, and applied with increasing percentages of ethanol in water (30, 50, 70, 95, 100) series to replace PBT. Samples were then immersed into xylene for clearance and mounted in DPX (Sigma, MI). The following primary antisera were used:

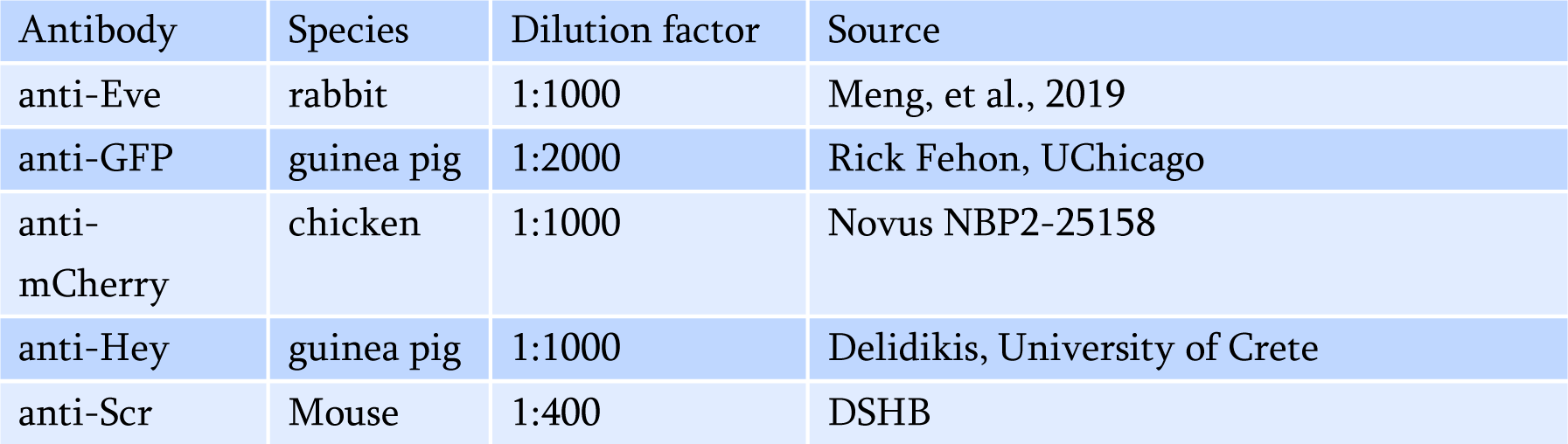

Secondary antibodies were obtained from ImmunoResearch (Bar Harbor, ME) and were used at a 1:400 dilution.

### Imaging, Statistics, Figure assembly

Images were acquired either on a Zeiss 800 confocal microscope with a 40X objective or a Zeiss 880 with Airyscan processing. Images were processed and analyzed using ImageJ (Fiji). Test and descriptive statistical analyses were performed using Prism (GraphPad). Figures were assembled in Adobe Illustrator.

### Cell counts

This procedure is for counting cells in a z-stack, particularly cells in a cluster that are challenging to count by eye. Using the Multi-point tool in FIJI, it is possible to label each cell with a numbered point to show the presence of all cells in a cluster. This works well for images taken at 2048 x 2048 with approx. 0.5 micron slices. (1) Open the .czi file in FIJI and select the channel of interest. (2) Go to Process/Enhance Contrast… and change Saturated Pixels to 0.1%. Select Normalize and Process all slices before pressing OK. (3) Go to Process/Noise/De-speckle to further process the image before analysis. (4) Save the changes as a .tif. (5) Open the .tif, scroll to the first slice where a cell appears, and zoom in on the cell cluster. Use the + and - keys to zoom to avoid moving points erroneously. (6) Select Multi-point in the FIJI toolbar and double-click to open the Point tool window. Set Type to Hybrid, Size to Tiny, and select Label points. Set counter to 1. Press OK. (7) With Multi-point tool selected, click on the center of a cell to drop a point. Scroll through the stack and drop point on each slice until the cell is no longer visible. With slices of approx. 0.5 microns cells should be present in 6-8 slices. (9) Return to the Point tool window and set counter to 2. Repeat to mark the remaining cells. (1) Notes specific to counting ELs: Label U neurons to ensure that Us are not counted as ELs, using the 0 counter.

### ts-MARCM experiments

We used an internal reference to precisely estimate when heat shocks were provided to the embryos to induce recombinase expression and initiate the lineage trace. This is because heat shocks were provided to embryos for 2-to 3-hour windows, it takes several hours for FLP to induce a genetic change (Wang et al. 2022), and neuroblasts divide every 45 minutes. This internal reference system works on the developmental principle that, unlike vertebrates and long-germ band insects, where axial development is temporally offset, with the anterior developing before the posterior, in short-germ band insects like *Drosophila,* segments develop synchronously. So, it is reasonable to assume that neuroblasts are dividing at approximately the same time along the A-P axis, and there are no differences between left and right hemi-segments. We found that most CNS had multiple clones (Figure 2E). For each CNS, we assessed each clone’s location and size. Because we know NB3-3 divides for the duration of embryogenesis (Gunnar et al. 2016), the number of EL labeled in the abdomen is directly related to the number of NB3-3 divisions yet to occur. If one or more clones were found within the abdomen, we used it to calculate the max abdominal clone size per CNS; using the max abdominal clone size serves as our internal reference. The maximum abdominal clone size is a proxy for the earliest an embryo could have been heat-shocked. We sorted each clone into bins based on its maximum abdominal clone size in its CNS. This binning generates the X-axis in Figure 2F and is labeled as “heat shock relative to inferred NB3-3 division number”.

