## Supplementary material for "How serially homologous neuroblasts produce different temporal cohorts along the *Drosophila* larval body axis": Figure

Supporting Information:

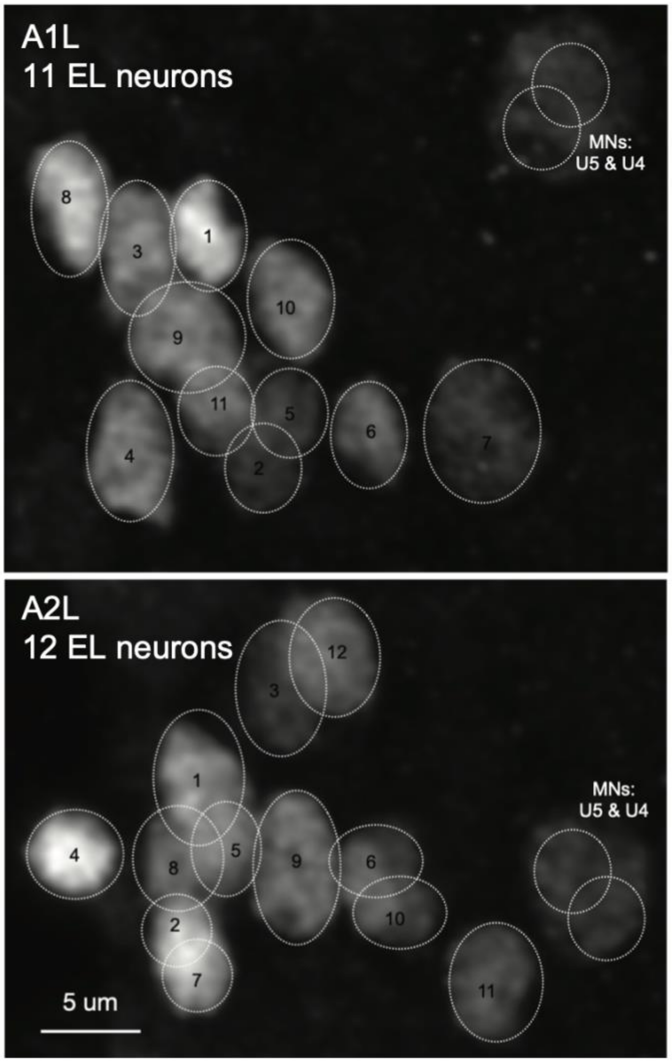


**Figure S1**. At the larval stage, there are 12 ELs in segment A2 compared to 11 in A1.

Images of laterally-located Eve(+) neurons in segments A1 and A2 of a larval CNS taken with high-resolution Airyscan microscopy. Images are Z-projections of the confocal stack stained with anti-Eve antibody. Each neuron is outlined by a circle. Two U motor neurons are located medially (right in these images). Each EL is numbered with lower numbers representing more neurons located more dorsally within the cluster. In segment A1, there are 11 EL neurons, and in segment A2, there are 12 EL neurons.

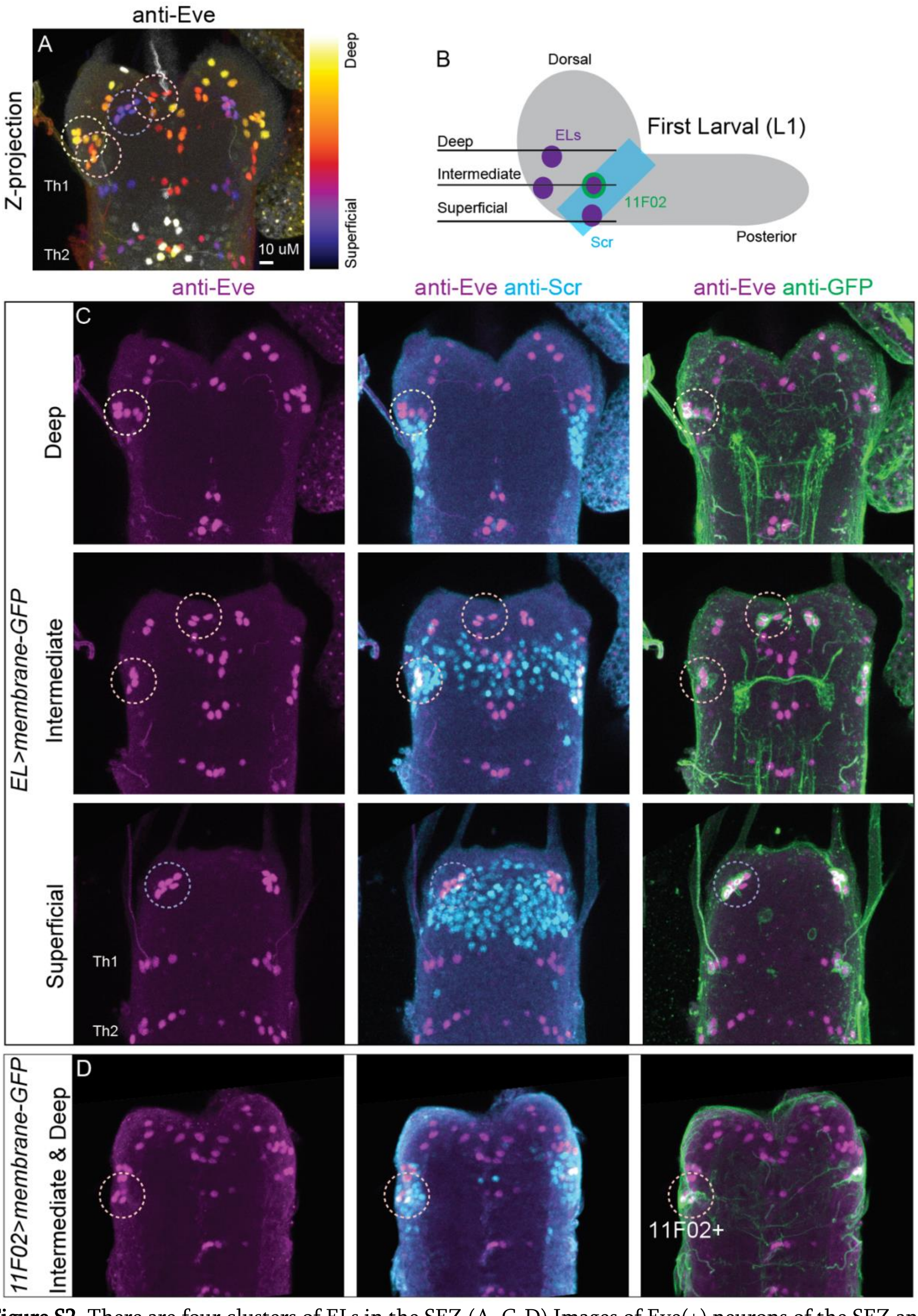


**Figure S2**. There are four clusters of ELs in the SEZ (A, C-D) Images of Eve(+) neurons of the SEZ and thorax. The dashed circles show the ELs on the left side of the SEZ.

(A) The image is a depth-coded Z-projection of anti-Eve staining. Warm colors (yellow) represent deeper slices, and cool (purple) colors represent superficial slices. Th1 and Th2 are the first two segments of the thorax. (B) Illustration of the CNS with approximate positions of EL clusters within the SEZ. (C-D) Eve is in magenta. Scr, in cyan, marks the SEZ-nerve cord border (Diao et al., 2024). Green shows anti-GFP staining. (C) The same CNS as in (A). GFP marks the ELs with *EL-GAL4>UAS-CD4-GFP*. (D) GFP marks 11F02(+) (late-born) neurons with *11F02-GAL4>UAS-CD4-GFP*. (C) Projections of the larval CNS's superficial, intermediate, and deep thirds. (D) Projection of the intermediate and deep thirds of the late-stage embryonic SEZ and thorax.

**Table S1**: Clones per CNS

See attached (Table too big to insert).

**Table S2**: Ts-MARCM dataset

| Heat shock protocol | Time of heat shock (minutes post fertilization) start | stop | CNSs | Clones number |
|---|---|---|---|---|
| a | 300 | 480 | 8 | 22 |
| b | 315 | 495 | 6 | 18 |
| c | 345 | 495 | 9 | 15 |
| d | 360 | 480 | 4 | 11 |
| e | 360 | 540 | 1 | 1 |
| f | 375 | 495 | 9 | 23 |
| g | 420 | 540 | 7 | 12 |
| h | 420 | 645 | 6 | 8 |
| j | 480 | 600 | 2 | 3 |
| i | 420 | 480 | 20 | 73 |
| k | 540 | 600 | 8 | 48 |

**Table S3**: NB3-3A neuron numbers are constant in nearly all segments.

| Segment | Stage 15 | Stage 17 | P-Value (ANOVA) |
|---|---|---|---|
| SEZ3 | 9.3+/-(n=4) | 11.8+/-0.5 (n=4) | 0.007 |
| Th1 | 5+/-0 (n=4) | 5+/-0 (n=4) | nd |
| Th2 | 5+/-0 (n=4) | 5+/-0 (n=4) | nd |
| Th3 | 5.8+/-0.5 (n=4) | 6+/-0 (n=4) | 0.40 |
| A1 | 1+/-0 (n=4) | 1+/-0 (n=4) | nd |
| A2 | 1+/-0 (n=4) | 1+/-0 (n=4) | nd |
| A3 | 1+/-0 (n=4) | 1+/-0 (n=4) | nd |
| A4 | 1+/-0 (n=4) | 1+/-0 (n=3) | nd |
| A5 | 1+/-0 (n=3) | 1+/-0 (n=4) | nd |
| A6 | 1+/-0 (n=4) | 1+/-0 (n=4) | nd |
| A7 | 1+/-0 (n=3) | 1+/-0 (n=3) | nd |
| Te1 | 1+/-0 (n=3) | 1+/-0 (n=3) | nd |
| Te2 | 0.5+/-0.7 (n=2) | 0.5+/-0.6 (n=4) | 1 |
| average +/- standard deviation (n=hemisegments counted) | | | |

**Table S4**: NB3-3B neurons lacking Eve expression increase in number in the terminus but are constant in other segments.

| Segment | Stage 15 | Stage 17 | P-Value (ANOVA) |
|---|---|---|---|
| SEZ3 | 0 +/- 0 (n=4) | 0 +/- 0 (n=4) | 1 |
| Th1 | 0 +/- 0 (n=4) | 0 +/- 0 (n=4) | 1 |
| Th2 | 0 +/- 0 (n=4) | 0 +/- 0 (n=4) | 1 |
| Th3 | 0 +/- 0 (n=4) | 0.3 +/- 0.6 (n=3) | 0.4 |
| A1 | 0.5 +/- 0.6 (n=4) | 0.5 +/- 0.6 (n=4) | 1 |
| A2 | 0 +/- 0 (n=4) | 0 +/- 0 (n=4) | 1 |
| A3 | 0 +/- 0 (n=4) | 0 +/- 0 (n=4) | 1 |
| A4 | 0 +/- 0 (n=4) | 0 +/- 0 (n=3) | 1 |
| A5 | 0 +/- 0 (n=3) | 0 +/- 0 (n=4) | 1 |
| A6 | 0 +/- 0 (n=4) | 0 +/- 0 (n=4) | 1 |
| A7 | 0.3 +/-0.5 (n=4) | 0.3 +/- 0.6 (n=3) | 1 |
| Te1 | 1.3 +/-1.2 (n=3) | 0.5 +/- 0.7 (n=2) | 0.4 |

| | | | |
|---|---|---|---|
| Te2 | 5.0 +/-0.0 (n=2) | 3.3 +/- 0.6 (n=3) | 0.04 |
| average +/- standard deviation (n=hemisegments counted) | | | |
